# PULPO: Pipeline of understanding large-scale patterns of oncogenomic signatures

**DOI:** 10.1101/2025.07.02.661487

**Authors:** Marta Portasany-Rodríguez, Gonzalo Soria-Alcaide, Elena G. Sánchez, María Ivanova, Ana Gómez, Reyes Jiménez, Jaanam Lalchandani, Gonzalo García-Aguilera, Silvia Alemán-Arteaga, Cristina Saiz-Ladera, Manuel Ramírez-Orellana, Jorge Garcia-Martinez

**Affiliations:** Fundación para la Investigación Biomédica Hospital Infantil Universitario Niño Jesús (FIBHNJS), Madrid, Spain; Hospital Infantil Universitario Niño Jesús (HNJS), Madrid, Spain; Instituto de Investigación Sanitaria La Princesa (IISLP), Madrid, Spain

## Abstract

**Summary:** PULPO v1.0 is a novel, fully automated pipeline designed for the preprocess and extraction of mutational signatures from raw Optical Genome Mapping (OGM) data. Built using Snakemake and executed within an isolated, Conda-managed environment, PULPO transforms complex cytogenetic alterations, captured at ultra-high resolution, into Catalogue of somatic mutations in Cancer (COSMIC)-based mutational signatures. This innovative approach not only enables researchers to work directly from raw OGM inputs but also streamlines the traditionally complex process of signature extraction, making advanced oncogenomic analyses accessible to users with varying levels of bioinformatics expertise. By facilitating the integration of comprehensive structural variants (SVs) and copy number variants (CNVs) data with established signature catalogs, PULPO paves the way for improved diagnostic accuracy and personalized therapeutic strategies.

**Availability and Implementation:** The pipeline is open source and freely available under the MIT License at https://github.com/OncologyHNJ/PULPO.

## 1. Introduction

SVs and CNVs are major classes of genomic alterations that play pivotal roles in carcinogenesis and tumor progression (Beroukhim et al. 2010; Weischenfeldt et al. 2013, Xu et al. 2022). Despite their biological significance, the systematic detection of these alterations has long been restricted by the limitations of conventional cytogenetic techniques such as karyotyping, fluorescence in situ hybridization (FISH), and multiplex ligation-dependent probe amplification (MLPA) (Park et al. 2011; Alías et al. 2012). These traditional methods often lack the resolution necessary to detect small CNVs or complex rearrangements, such as chromothripsis or balanced translocations, and are frequently hampered by high costs and labor-intensive workflows.

The recent advent of OGM has transformed the landscape of genomic analysis by enabling genome-wide detection of SVs and CNVs at an ultra-high resolution (<1 kb) in a single assay (Levy et al. 2024; Neveling et al. 2021). Moreover, OGM is capable of detecting a wide range of chromosomal alterations across the entire genome, not limited to restricted areas. Studies have demonstrated that OGM offers superior sensitivity and resolution compared to standard cytogenetic methods, reporting up to a 30 % increase in diagnostic yield for both hematologic malignancies and solid tumors (Neveling et al. 2021; Lühmann et al. 2023). This breakthrough technology not only refines the detection of structural variants but also provides an unprecedented opportunity to explore the intricate molecular architecture that underpins tumorigenesis.

Beyond the detection of genomic alterations, the analysis of mutational signatures—defined as the patterns of somatic alterations left by various endogenous processes (e.g., defective DNA repair mechanisms) or exogenous exposures (e.g., tobacco smoking and UV radiation)—has emerged as a crucial tool for understanding cancer etiology (Steele et al. 2022). While the mutational landscapes of single-base substitutions (SBS), doublet base substitutions (DBS), and small insertions/deletions (ID) have been extensively catalogued in resources like COSMIC (Alexandrov et al. 2020), structural mutational signatures associated with SVs and CNVs remain less characterized. The limited resolution of prior genomic platforms has prevented a systematic interrogation of the complex breakpoint patterns that could unveil novel insights into the mutagenic processes driving cancer (Degasperi et al. 2020; Cortés-Ciriano et al. 2023, Dentro et al. 2021).

The integration of OGM with mutational signature analysis represents a highly promising frontier in precision oncology. SigProfiler, widely recognized for its robust performance in identifying similarities with COSMIC signatures, has firmly established itself as a powerful tool in this domain (Islam et al. 2020; Islam et al. 2022). However, its implementation necessitates specialized bioinformatics expertise and programming skills. To broaden access to mutational signature analysis and enhance usability, we have developed PULPO (Pipeline of Understanding Large-scale Patterns of Oncogenomic signatures). PULPO incorporates the similarity search capabilities of SigProfiler within an intuitive, user-friendly framework, while also pioneering the use of OGM data. This innovation not only streamlines the analytical workflow but also offers a novel perspective for elucidating the mutagenic processes that underlie cancer development.

PULPO distinguishes itself by integrating high-resolution SV/CNV data obtained through OGM with the comprehensive mutational signature repository provided by COSMIC (Alexandrov et al. 2020). Utilizing similarity-based algorithms, the pipeline systematically compares OGM-derived variant profiles to the COSMIC catalog, enabling the precise identification of structural signatures. Built on a preconfigured Snakemake pipeline, PULPO streamlines data preprocessing, signature similarity scoring, and visualization, thereby democratizing access to this advanced analysis even for researchers with limited bioinformatics expertise.

The clinical implications of PULPO are substantial. By linking structural mutational signatures to specific etiological processes and therapeutic vulnerabilities, PULPO not only enhances our understanding of the molecular drivers of cancer but also opens new avenues for translational research. For instance, matching signature profiles to therapy-associated patterns such as those related to APOBEC activity or homologous recombination deficiency can facilitate drug repositioning and the identification of actionable targets (Alexandrov et al. 2020, Watkins et al. 2023). Additionally, the detection of exogenous exposure signatures such as tobacco-associated clustered translocations can guide preventive strategies for high-risk patient populations (Alexandrov et al., 2016). Furthermore, recent work has demonstrated that geographic variation of mutagenic exposures can significantly impact cancer genomes (Senkin et al. 2024). In summary, by leveraging the transformative resolution of OGM, PULPO bridges a critical gap in structural mutational signature analysis and holds significant promise for clinical applications, ranging from improved diagnostic accuracy to the personalization of cancer treatment.

## 2. Methods

PULPO v1.0 was developed using Snakemake v7.32.4, a robust workflow management system that ensures scalability, reproducibility, and modularity in bioinformatics pipelines (Mölder et al. 2021). Its design strongly emphasizes automation and user transparency, enabling integration into high-throughput computational environments. To ensure reproducibility across platforms and users, PULPO is executed within a fully controlled and isolated Conda environment (Anaconda 2016), which includes all the necessary dependencies and packages specified in an environment file. The pipeline workflow is summarized in the graphical abstract, and the main steps of the analysis are detailed below.

### Validation and Pre-processing of Input Files

Before initiating any downstream analysis, PULPO performs a rigorous quality control step by validating input files to ensure that no sample is empty. If an empty sample file is detected, the pipeline halts execution immediately and generates an informative error log, automatically stored in the designated log directory within the user-specified working folder.

Pre-processing of data includes a dedicated module for raw OGM data generated by Bionano Access v1.6.1 and v1.8.1. This step addresses a current gap in standardized workflows for extracting mutational signatures from OGM, which remains a novel and underdeveloped data source in this context.

The data conversion module takes raw OGM data and transforms it into a format suitable for the SigProfiler tool suite. Specifically, it converts OGM-derived CNV data into BED format and SV data into BEDPE format, both widely used and standardized formats in numerous bioinformatics tools. This not only enables their use within the PULPO framework but also facilitates their integration with any other bioinformatics tool that requires these file types. Since OGM files are converted into BED and BEDPE formats, the tool can alternatively be initialized directly with these standard file types.

Once the data are formatted, they are processed using SigProfilerMatrixGenerator v1.2.28 and SigProfilerExtractor v1.1.24 (Islam et al. 2020; Islam et al. 2022). The pipeline offers two main modes of analysis:

1. Per-sample analysis for individual-level signature exploration.
2. Combined cohort + sample analysis to facilitate broader comparative analyses while preserving individual resolution.

By default, PULPO employs the GRCh38 human reference genome to ensure consistency; however, its modular architecture allows future extension to other reference builds, based on user requirements. To enhance usability and flexibility, several key parameters of the SigProfilerExtractor tool, such as *minimum_signatures, maximum_signatures*, and *nmf_replicates*, can be explicitly defined by the user or selected automatically using optimized, dataset-specific default settings, thereby lowering the entry barrier for less experienced users while permitting fine-tuning for advanced applications.

Importantly, PULPO enables the analysis of SVs and CNVs either independently or in combination, offering a versatile and customizable approach for researchers interested in exploring the full spectrum of large-scale genomic rearrangements and their associated mutational processes. This makes the tool particularly valuable in cancer genomics studies, where such alterations play a key role.

## 3. Availability and Implementation

PULPO is open-source, actively maintained, and freely available under the MIT License. The full source code, documentation, and example datasets can be accessed through its GitHub repository: https://github.com/OncologyHNJ/PULPO.

The pipeline is designed for Unix-based systems (Linux/macOS) and can be executed locally or within high-performance computing environments. Although not natively compatible with Windows, PULPO could be run on Windows systems using Docker, which allows encapsulation of the full environment and dependencies, thereby ensuring reproducibility and ease of deployment.

After cloning the repository, users can customize the execution by editing the config.yml file, which serves as the central configuration for specifying input paths, analysis modes, and key parameters. A preconfigured example is provided to facilitate initial runs and to adapt the pipeline to diverse use cases. To run PULPO, users must have a functional Conda installation (e.g., via Miniconda) and Snakemake. Detailed installation and execution instructions are included in the repository, which also provides guidance on setting up the environment and executing the workflow with minimal effort. PULPO supports parallel execution and is suitable for large-scale genomic datasets.

## 4. Results

To evaluate the feasibility and robustness of PULPO, we conducted a proof-of-concept analysis on 130 paediatric samples of patients diagnosed with acute lymphoblastic leukaemia (ALL). The samples were profiled using OGM technology (Bionano Genomics) in conjunction with the Rare Variant Analysis pipeline. The use of ALL patient samples has been approved by the CEIm at Hospital Infantil Universitario Niño Jesús (HUNJ) under internal code R-0017/13 (approved on May 8, 2013), and written informed consent was obtained from all participants or their legal guardians, in accordance with the Declaration of Helsinki and applicable national regulations on biomedical research involving human subjects.

PULPO was executed on a standard computer equipped with 16 GB RAM, an 11th Gen Intel® Core™ i7-1165G7 @ 2.80GHz × 8 CPU, and Ubuntu 22.04.5 LTS, thereby demonstrating its compatibility with widely accessible computing infrastructures.

The pipeline successfully processed the majority of samples by automatically executing all the required steps for mutational signature extraction, generating standard outputs which include signature decomposition plots, reconstruction metrics, and comprehensive quality control summaries.

For most of the 130 samples analyzed, the pipeline generated valid mutational signature profiles. In a minority of cases, however, no signatures could be extracted due to insufficient input complexity. Given that the COSMIC signature framework is based on comparing the relative contributions of multiple mutational processes, an insufficient number of genomic events prevent robust decomposition and reliable signature assignment. The capacity of PULPO to automatically identify and exclude such samples ensures analytical rigor and output reliability.

To demonstrate PULPO’s ability to extract biologically meaningful mutational signatures from OGM data, we applied the pipeline to the 130-sample paediatric ALL cohort. PULPO assigned a median of 3 COSMIC signatures per sample (range 1–5), with the most prevalent signatures being SV2 (associated with non-clustered translocations), SV7 (non-clustered deletions), CN2 (ploidy gain (1x) / aneuploidy), CN4 (chromotripsis) and CN25 (MMR deficiency) (Supplementary Fig. S1). The mean cosine similarity between each sample’s to known COSMIC structural signatures observed variant spectrum and its reconstructed signature profile was 0.88 (range: 0.88–0.96), indicating high fidelity of signature decomposition.

The mean cosine similarity between observed and reconstructed structural variant profiles across samples, based on known COSMIC signatures, was 0.88 (range: 0.88–0.96), indicating accurate signature attribution (Supplementary Tables S1 and S2). Cosine similarity values below 0.8 are considered insufficient for confident assignment to know COSMIC structural signatures (observed in some cases, see Supplementary Table S2). Overall, these values support a high fidelity of signature decomposition. Together, these results demonstrate that PULPO effectively automates COSMIC signature assignment on OGM data and captures mutational profiles consistent with known structural variant etiologies.

Currently, there is no established benchmark or reference pipeline for the extraction of COSMIC-like signatures from OGM-derived data, and to the best of our knowledge, PULPO is the first tool specifically designed to bridge this gap. Although the COSMIC mutational signature catalog was not originally developed using OGM data, we posit that the high resolution and structural accuracy of OGM render it a compelling source for signature inference, especially for detecting large-scale genomic rearrangements that are often underrepresented in short-read sequencing data.

## 5. Discussion and Conclusion

In this study, we demonstrate that PULPO enables a robust and accessible framework for structural variant signature analysis by leveraging high-resolution SV/CNV data generated via OGM. By systematically integrating these profiles with the COSMIC repository of mutational signatures (Alexandrov et al. 2020) through similarity-based algorithms, PULPO facilitates the accurate identification of structural mutational patterns across samples. Its implementation within a preconfigured Snakemake pipeline not only ensures reproducibility and scalability, but also significantly lowers the barrier to entry for researchers without extensive bioinformatics training. This integrative approach highlights the potential of OGM-based signature analysis as a complementary tool for genomic characterization, particularly in settings where conventional sequencing may overlook complex structural rearrangements.

PULPO presents significant potential for clinical translation by enabling the interpretation of structural mutational signatures in the context of underlying etiological mechanisms. Through this approach, it contributes to a deeper understanding of the molecular processes driving tumorigenesis and supports the development of precision oncology strategies. Notably, the ability to associate specific signature profiles with therapeutic contexts—such as APOBEC activity or homologous recombination deficiency (HRD)—may inform drug repurposing efforts and highlight novel therapeutic vulnerabilities (Alexandrov et al. 2020; Watkins et al. 2023). Moreover, the identification of mutational patterns linked to environmental or lifestyle exposures, including tobacco-related clustered translocations, could aid in stratifying at-risk populations and designing targeted prevention programs (Alexandrov et al. 2016). Together, these applications underscore PULPO’s potential to serve as a bridge between structural variant research and actionable clinical insights.

Looking forward, as OGM becomes more widely adopted and the volume of available datasets increases, we anticipate that de novo mutational signatures will be reconstructed directly from OGM-based cohorts. PULPO was developed with this future evolution in mind; its modular structure enables integration of updated COSMIC releases or alternative, OGM-specific signature catalogs with minimal adjustments, ensuring long-term utility and adaptability. Recent work employing bioinformatic methods to identify mutational signatures further supports the evolution of such approaches (Islam & Alexandrov 2021). This innovative integration not only enhances our understanding of the complex mutagenic processes driving cancer but also holds significant promise for clinical applications, including improved diagnostic accuracy, refined therapeutic stratification, and personalized treatment approaches.

## Acknowledgements

The authors sincerely appreciate the work of the whole FIBHNJS staff.

## Author contributions

Marta Portasany-Rodríguez (Conceptualization [equal], Data curation [equal], Formal analysis [lead], Investigation [equal], Methodology [equal], Project administration [equal], Resources [equal],

Software [equal], Validation [equal], Visualization [equal], Writing – original draft [equal], Writing –review & editing [equal])

Gonzalo Soria-Alcaide (Conceptualization [equal], Software [equal], Supervision [equal], Writing – review & editing [equal])

Elena G. Sánchez (Processing of ALL pediatric samples by OGM [lead], Sample sequencing [lead], Pre-analysis of results [lead], Writing – review & editing [equal])

María Ivanova (Processing of ALL pediatric samples by OGM [equal])

Ana Gómez (Processing of ALL pediatric samples by OGM [equal])

Reyes Jiménez (Processing of ALL pediatric samples by OGM [equal])

Jaanam Lalchandani (Writing – review & editing [equal])

Gonzalo García-Aguilera (Writing – review & editing [equal])

Silvia Alemán-Arteaga (Writing – review & editing [equal])

Cristina Saiz-Ladera (Writing – review & editing [equal])

Manuel Ramírez-Orellana (Conceptualization [equal], Project administration [equal], Resources [equal], Supervision [lead], Writing – review & editing [equal])

Jorge Garcia-Martínez (Conceptualization [equal], Data curation [equal], Formal analysis [equal], Investigation [equal], Methodology [equal], Project administration [equal], Resources [equal], Software [equal], Supervision [lead], Validation [equal], Visualization [equal], Writing – original draft [equal], Writing – review & editing [equal])

## Graphical abstract

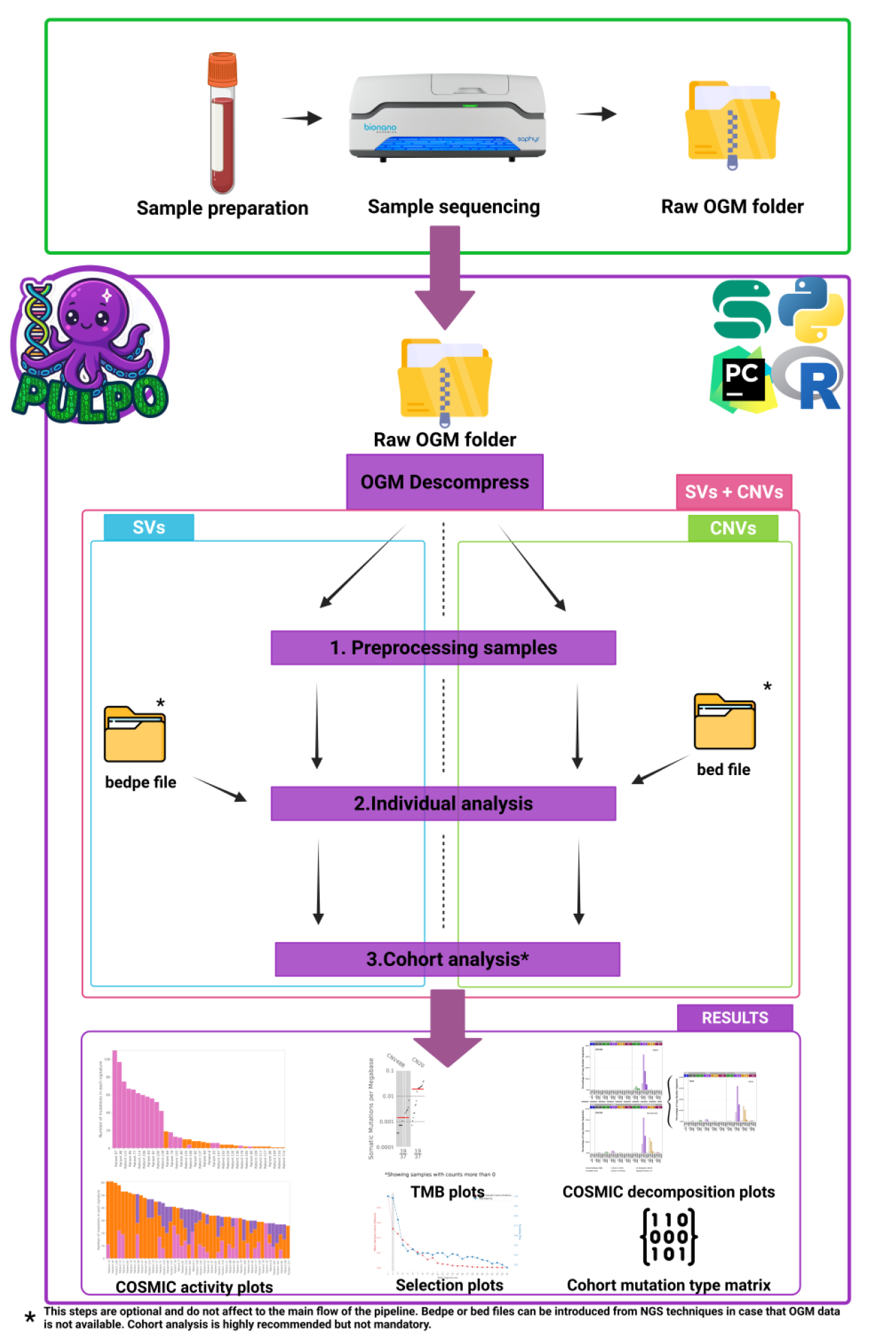

Schematic overview of the PULPO workflow. Starting from raw OGM data, users can perform SVs-only (left), CNVs-only (right), or combined SVs+CNVs (left and right) analyses, as indicated in purple. The workflow generates standard

SigProfiler outputs, including COSMIC signature activity plots, tumor mutational burden (TMB) plots, selection plots, and COSMIC decomposition plots, along with mutation type matrices at both cohort and individual levels.

## References

1. Alías L, Bernal S, Barceló MJ, Also-Rallo E, Martínez-Hernández R, Rodríguez-Alvarez FJ, et al. Accuracy of marker analysis, quantitative real-time polymerase chain reaction, and multiple ligation-dependent probe amplification to determine SMN2 copy number in patients with spinal muscular atrophy. Genet Test Mol Biomarkers. 2011;15(9):587–594. 10.1089/gtmb.2010.0253.

2. Alexandrov LB, Ju YS, Haase K, Van Loo P, Martincorena I, Nik-Zainal S, et al. Mutational signatures associated with tobacco smoking in human cancer. Science. 2016;354(6312):618–622. 10.1126/science.aag0299.

3. Alexandrov LB, Kim J, Haradhvala NJ, Huang MN, Ng AWT, Wu Y, et al. The repertoire of mutational signatures in human cancer. Nature. 2020;578(7793):94–101. 10.1038/s41586-020-1943-3.

4. Anaconda, Inc. (2016) Conda: Package, dependency and environment management. Available at: https://docs.conda.io

5. Beroukhim R, Mermel CH, Porter D, Wei G, Raychaudhuri S, Donovan J, et al. The landscape of somatic copy-number alteration across human cancers. Nature. 2010;463(7283):899–905. 10.1038/nature08822.

6. Cortés-Ciriano I, Lee JJK, Xi R, Jain D, Jung YL, Yang L, et al. Comprehensive analysis of chromothripsis in 2,658 human cancers using whole-genome sequencing. Nat Genet. 2020;52(3):331–341. 10.1038/s41588-019-0576-7.

7. Degasperi A, Zou X, Amarante TD, et al. A practical framework and online tool for mutational signature analyses show inter-tissue variation and driver dependencies. Nat Cancer. 2020;1(2):249– 263. 10.1038/s43018-020-0027-5.

8. Dentro SC, Leshchiner I, Haase K, Tarabichi M, Wintersinger J, Deshwar AG, et al. Characterizing genetic intra-tumor heterogeneity across 2,658 human cancer genomes. Cell. 2021;184(8):2239– 2254.e39. 10.1016/j.cell.2021.03.009.

9. Islam SMA, Alexandrov LB. Bioinformatic methods to identify mutational signatures in cancer. In: Cobaleda C, Sánchez-García I, eds. Leukemia Stem Cells: Methods and Protocols. Springer US; 2021:447–473.

10. Islam SMA, Díaz-Gay M, Wu Y, et al. SigProfilerMatrixGenerator: a tool for visualizing and exploring patterns of small mutational events. BMC Genomics. 2020;21(1):259. 10.1186/s12864-020-6723-6.

11. Islam SMA, Díaz-Gay M, Wu Y, et al. Uncovering novel mutational signatures by de novo extraction with SigProfilerExtractor. Cell Genom. 2022;2(11):100179. 10.1016/j.xgen.2022.100179.

12. Levy B, Kanagal-Shamanna R, Sahajpal NS, Neveling K, Rack K, Dewaele B, et al. A framework for the clinical implementation of optical genome mapping in hematologic malignancies. Am J Hematol. 2024;99(4):642–661. 10.1002/ajh.27175.

13. Mölder F, Jablonski KP, Letcher B, Hall MB, Tomkins-Tinch CH, Sochat V, et al. Sustainable data analysis with Snakemake. F1000Res. 2021;10:33. 10.12688/f1000research.29032.2.

14. Neveling K, Mantere T, Vermeulen S, Oorsprong M, van Beek R, Kater-Baats E, et al. Next-generation cytogenetics: Comprehensive assessment of 52 hematological malignancy genomes by optical genome mapping. Am J Hum Genet. 2021;108(8):1423–1435. 10.1016/j.ajhg.2021.06.001.

15. Park SJ, Jung EH, Ryu RS, et al. Clinical implementation of whole-genome array CGH as a first-tier test in 5080 pre and postnatal cases. Mol Cytogenet. 2011;4:12. 10.1186/1755-8166-4-12.

16. Senkin S, Moody S, Díaz-Gay M, Abedi-Ardekani B, Cattiaux T, Ferreiro-Iglesias A, et al. Geographic variation of mutagenic exposures in kidney cancer genomes. Nature. 2024;629(8013):910–918. 10.1038/s41586-024-07368-2.

17. Steele CD, Abbasi A, Islam SMA, Bowes AL, Khandekar A, Haase K, et al. Signatures of copy number alterations in human cancer. Nature. 2022;606(7916):984–991. 10.1038/s41586-022-04738-6.

18. Watkins JA, Jonsson P, Van Hoeck A, et al. Homologous recombination deficiency signatures predict PARP inhibitor sensitivity across cancer types. Cancer Discov. 2023;13(5):1216–1233. 10.1158/2159-8290.CD-22-1336.

19. Weischenfeldt J, Symmons O, Spitz F, Korbel JO. Phenotypic impact of genomic structural variation: insights from and for human disease. Nat Rev Genet. 2013;14(2):125–138. 10.1038/nrg3373.

20. Xu Z, Lee DS, Chandran S, Le VT, Bump R, Yasis J, et al. Structural variants drive context-dependent oncogene activation in cancer. Nature. 2022;612(7940):564–572. 10.1038/s41586-022-05504-4.

