## Supplementary Information for "PULPO: Pipeline of understanding large-scale patterns of oncogenomic signatures"

***Figures***

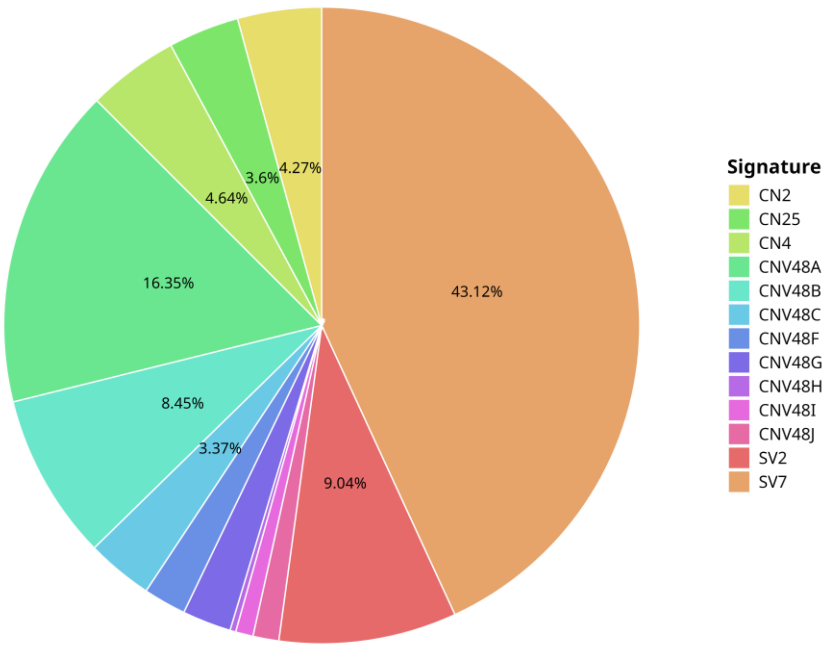


**Figure S1:**

Schematic overview of PULPO results. Relative contribution of each mutational signature activity in the cohort including CN and SV signatures. Signature activity is represented as the total number of mutations attributed to each signature.

***Tables***

| Signature | Percent |
|---|---|
| SV7 | 97.5 % (118/121) |
| SV2 | 41.3 % (50/121) |
| SV2 + SV7 | 38.8 % (47/121) |

**Table S1:**

Proportion of SV signatures in the cohort. In parentheses, the number of samples in which each signature is present in the cohort.

| Signature | Percent |
|---|---|
| CNV48C | 55.2 % (64/116) |
| CNV48A | 50.9 % (59/116) |
| CNV48B | 46.6 % (54/116) |
| CNV48A + CNV48B | 40.5 % (47/116) |
| CN4 | 31 % (36/116) |
| CN25 | 24.1 % (28/116) |
| CNV48F | 23.3% (27/116) |
| CNV48J | 22.4% (26/116) |
| CNV48G | 21.6% (25/116) |
| CNV48A + CNV48C | 21.6% (25/116) |
| CN2 | 19% (22/116) |
| CN2 + CN25 | 2.6 % (3/116) |
| CN2 + CN4 | 1.7 % (2/116) |

**Table S2:**

Selected proportions of CNV signatures and their combinations in the cohort, ordered by percentage. In parentheses, the number of samples in which each mutational signature is present. Signatures starting with "CN" correspond to the COSMIC collection. Signatures starting with "CNV" are cohort-specific, with a cosine similarity < 0.8 to known COSMIC signatures, indicating no direct decomposition into the COSMIC signature collection.
